# Identification and characterization of novel mutants of Nsp13 protein among Indian SARS-CoV-2 isolates

**DOI:** 10.1101/2021.07.30.454406

**Authors:** Deepa Kumari, Namrata Kumari, Sudhir Kumar, Prabhat Kumar Sinha, Shivendra Kumar Shahi, Nihar Ranjan Biswas, Abhay Kumar

## Abstract

SARS-CoV-2, the causative agent of COVID-19 has mutated rapidly which enabled them to adapt and evade the immune system of the host. Emerging SARS-CoV-2 variants with crucial mutations pose a global challenge in context of therapeutic drugs and vaccines being developed globally. There are currently no specific therapeutics or vaccines available to combat SARS-CoV-2 devastation. In view of this, the current study aimed to identify and characterize the mutations found in the Nsp13 of SARS-CoV-2 in Indian isolates. Non-structural protein, Nsp13 protein sequences from Indian isolates were analyzed by comparing with the first reported Severe acute respiratory syndrome Corona Virus-2 (SARS-CoV-2) protein sequence from Wuhan, China. Out of 825 Nsp13 protein sequences, a total of 38 mutations were observed among Indian isolates. Our data show that mutations in Nsp13 at various positions (H164Y, A237T, T214I, C309Y, S236I, P419S, V305E, G54S, H290Y, P53S, A308Y, and A308Y) have a significant impact on the protein’s stability and flexibility. Also, the impact of Nsp13 mutations on the protein function were predicted based on PROVEAN score that includes 15 mutants as neutral and 23 mutants as deleterious effect. Furthermore, B-cell epitopes contributed by Nsp13 were identified using various predictive immunoinformatic tools. Immunological Parameters of Nsp13 such as antigenicity, allergenicity and toxicity were evaluated to predict the potential B-cell epitopes. The predicted peptide sequences were correlated with the observed mutants. Our predicted data showed that there are seven high rank linear epitopes as well as 18 discontinuous B-cell epitopes based on immunoinformatic tools. Moreover, it was observed that out of total 38 identified mutations among Indian SARS-CoV-2 Nsp13 protein, four mutant residues at position 142 (E142), 245 (H245), 247 (V247) and 419 (P419) are localised in the predicted B cell epitopic region. Altogether, the results of the present *in-silico* study might help to understand the impact of the identified mutations in Nsp13 protein on its stability, flexibility and function.

## INTRODUCTION

A recent emergence with an outburst of coronavirus disease 2019 (COVID-19) pandemic caused by novel beta coronavirus, severe acute respiratory syndrome coronavirus 2 (SARS-CoV-2) [1] identified first in Wuhan, China, in late Dec 2019 [2-5]. COVID-19 spread rapidly across the worldwide population and was declared as a global pandemic on 11 March 2020 by World Health Organization (WHO). It has infected hundreds of millions of people and killed more than three million people worldwide. The world is in the midst of second wave of pandemic with rapidly evolving new variants of SARS-CoV-2 resulting in surge of the COVID-19 cases worldwide. The highly contagious nature of SARS-CoV-2 has now developed into a severe threat to global public health. The manifestations of SARS-CoV-2 ranges from asymptomatic, common cold to lethal viral respiratory illness in infected individuals [6].

Coronaviruses are enveloped with a non-fragmented, large single stranded (positive sense) RNA virus with GC content ranging from 32% to 43%. They are classified into four genera, namely, alphacoronavirus (α CoV), Beta coronavirus (β CoV), gamma coronavirus (γ CoV) and delta coronavirus (δ CoV), respectively [7]. SARS-COV-2 belongs to β coronavirus 2b lineage with a genome of length 29.9 kb which encodes 29 proteins. [8]. The genomic sequence of novel CoV, SARS-COV-2 exhibits similarity to 79.6% with SARS-CoV and about 50% with MERS-CoV (Middle East respiratory syndrome coronavirus), respectively.

The phylogenetic analysis has illustrated that SARS-CoV-2 is relatively more similar to SARS-CoV than MERS-CoV. Based on homology modelling studies, SARS-CoV-2 shares 96.2% nucleotide similarity with RaTG13, a bat CoV from *Rhinolophus Affinis*. The genomic organisation of SARS-CoV-2 includes ‘5 UTR, ORF1ab, S, E, M, N and 3’UTR. Its genome comprises of ORFs which encode for four major structural proteins, Spike glycoprotein (S), Envelope (E), membrane (M) and Nucleocapsid (N) proteins), 16 Nsps (non-structural proteins) and nine other accessory proteins [9-11]. ORF1a/b region is the largest ORF that covers about two-third of whole genome length which encode known sixteen non-structural proteins (16Nsps). ORFs in the remaining one third of the genome near the 3’ end codes for major structural and other accessory proteins. The ribosomal frameshift (−1) between ORF1a and ORF1b produces two polypeptides (pp1a and pp1ab) which are proteolytically cleaved by viral proteases, namely, main protease (M pro) or Chymotrypsin-like protease (3CLpro) and papain-like protease (PL pro) to release 16 Nsps of 7096 amino acids length. These Nsps are crucially important in viral replication and transcription processes [8]. Nsps are considered more conserved than SARS-CoV-2 structural protein [12]. Among the 16 known SARS-CoV-2 Nsp proteins, the Nsp13 helicase is a crucial component for viral replication [13] and the most conserved non-structural protein with 99.8% sequence identity to SARS-CoV Nsp13 [14]. The highest sequence conservation shared by SARS-CoV-2 Nsp13 within the CoV family, reflects their significant role in viral viability. The Nsp13, RNA helicase enzyme, also represents a promising therapeutic target for anti-CoV drug development [15-18].

The ongoing rapid transmission and global spread of SARS-CoV-2 has acquired various mutations. The global advancement in the sequencing efforts of SARS-CoV-2 have reported several mutations in their proteins like spike protein, nucleocapsid, PLpro and ORF3a from different population [19-21]. Emerging variants of SARS-CoV-2 with crucial mutations is a sparkling challenge in context of therapeutics drug and vaccines developed globally. Till date, there is no specific therapeutics and vaccines available to combat SARS-CoV-2. In view of this, the present study is focussed on identifying and characterizing the mutations in SARS-CoV-2 Nsp13 protein that might prove to be useful towards the development of drugs and effective vaccines against this highly mutable coronavirus. It could also help in improving the current diagnostic approaches for viral detection, thereby controlling the transmission of virus. As a consequence of the indispensable role of SARS-Cov-2 Nsp13 in the replication of virus, the study is aimed to understand the possible structural and functional implications of Nsp13 mutations among Indian isolates.

## MATERIAL AND METHODS

### Sequence retrieval and multiple sequence alignments

As of 24 May, 2021, there were 825 ORF1ab protein sequences of Indian SARS-CoV-2 deposited in NCBI Virus database. All these ORF1ab protein sequences (7096 residues) comprising all non-structural proteins (Nsp1-16) were retrieved from the NCBI-Virus database. The protein accession numbers of ORF1ab Sequences used in the study are mentioned in the supplementary Table1. The first reported sequence from China was used as reference or wild type ORF1ab sequence (accession number: YP_009724389) for mutational analysis in the study. The polypeptide sequences of the ORF1ab were exported in the FASTA format in which the Nsp13 of 601 residues starting from 5325 to 5925 were extracted from the ORF1ab sequence. Nsp13 is proteolytically cleaved by the protease encoded by the SARS-CoV-2 genome. The Clustal omega online webserver tool was used for multiple sequence alignments (MSA) [22,23] to identify the mutations in the amino acid sequences of Nsp13 among the Indian isolates. Clustal omega is widely used online program for generating alignments efficiently using fast and reliable algorithm. For this analysis, the Indian sequences were compared with the reference sequence of Nsp13 (accession number: YP_009724389) and then the identified mutant residues were marked carefully and noted for further analysis.

**Table 1.** list of identified mutations in Nsp13 among Indian isolates.

| S. No. | Mutation | Polarity changes | Charge changes |
| --- | --- | --- | --- |
| 1 | P53S | NP to P | Neutral to Neutral |
| 2 | G54S | P to P | Neutral to Neutral |
| 3 | T58A | P to NP | Neutral to Neutral |
| 4 | M68K | NP to P | Neutral to Basic |
| 5 | V98F | NP to NP | Neutral to Neutral |
| 6 | S100G | P to P | Neutral to Neutral |
| 7 | E142V | P to NP | Acidic to Neutral |
| 8 | V154L | NP to NP | Neutral to Neutral |
| 9 | H164Y | P to P | Basic to Neutral |
| 10 | S166A | P to NP | Neutral to Neutral |
| 11 | T214I | P to NP | Neutral to Neutral |
| 12 | V226L | NP to NP | Neutral to Neutral |
| 13 | L227Q | NP to P | Neutral to Neutral |
| 14 | T228K | P to P | Neutral to Basic |
| 15 | S236I | P to NP | Neutral to Neutral |
| 16 | A237T | NP to P | Neutral to Neutral |
| 17 | H245R | P to P | Basic to Basic |
| 18 | V247F | NP to NP | Neutral to Neutral |
| 19 | Y253H | P to P | Neutral to Basic |
| 20 | S259T | P to P | Neutral to Neutral |
| 21 | M274V | NP to NP | Neutral to Neutral |
| 22 | H290Y | P to P | Basic to Neutral |
| 23 | A296S | NP to P | Neutral to Neutral |
| 24 | L297F | NP to NP | Neutral to Neutral |
| 25 | P300L | NP to NP | Neutral to Neutral |
| 26 | I304K | NP to P | Neutral to Basic |
| 27 | V305E | NP to P | Neutral to Acidic |
| 28 | A308Y | NP to P | Neutral to Neutral |
| 29 | C309Y | P to P | Neutral to Neutral |
| 30 | S310Y | P to P | Neutral to Neutral |
| 31 | D315E | P to P | Acidic to Acidic |
| 32 | L384M | NP to NP | Neutral to Neutral |
| 33 | R392C | P to P | Basic to Neutral |
| 34 | P419S | NP to P | Neutral to Neutral |
| 35 | R442Q | P to P | Basic to Neutral |
| 36 | P504S | NP to P | Neutral to Neutral |
| 37 | P504L | NP to NP | Neutral to Neutral |
| 38 | I575L | NP to NP | Neutral to Neutral |

### Prediction of effect of mutations on protein stability and dynamicity

In order to understand the impact of mutational changes on the protein stability and dynamicity of Nsp13, protein modelling studies were performed by using DynaMut, a webserver tool [24]. The 3D structure of target protein, Nsp13 with PDB ID-6ZSLwas used for protein modelling studies. DynaMut programme is widely used to analyse and visualise the changes in protein dynamics and stability caused by mutations in the protein in terms of difference in vibrational entropy (ΔΔS) and free energy(ΔΔG) between wild type and mutant protein. The effect of mutations on change in protein stability was estimated by predicted ΔΔG(Kcal/mol) between wild type and mutant protein. The positive values (more than zero) indicate the stabilization of protein structure and negative values (below zero) show the destabilization of protein structure. The Protein flexibility and rigidity were calculated by difference in vibrational entropy (ΔΔS) between wild type and mutant protein of Nsp13. The positive ΔΔS values correspond to increase in the flexibility of Nsp13 protein whereas, the negative ΔΔS values represent the decrease in flexibility of protein.

### Prediction of PROVEAN Score

To understand the effect of amino acid variations on the function of Nsp13, PROVEAN (Protein Variation effect Analyzer) tool was used [25]. This prediction tool was used to generate a PROVEAN Score for each variant. PROVEAN provides an approach to predict the functionally important sequence variants. Based on predicted PROVEAN Score, mutants are classified as deleterious or neutral. The protein variant is predicted to have a deleterious effect, if the PROVEAN Score is equal to or less than the default threshold score (−2.5) and if PROVEAN Score is more than the threshold score the variant is predicted to have a neutral effect. PROVEAN tool is useful to measure the functional effect of protein sequence variations including amino acid substitutions, insertions, and deletions.

### Predictions of B-cell epitopes for Nsp13

B-cell epitopes predictions were performed using an online webserver, IEDB (Immune Epitope Database) prediction tool [26] as described by Jesperson et al [27]. This prediction tool predicts the linear continuous B-Cell epitopes of Nsp13 based on Bepipred linear epitope prediction method using all the standard parameters such as flexibility, hydrophilicity, accessibility and beta turns at a threshold value of 0.48. The identified epitopes of Nsp13 were further assessed for immunological parameters such as antigenicity, allergenicity and toxicity (by Toxin Pred). Antigenicity and Allergenicity of peptides were predicted by a freely available webserver, VaxiJen (http://www.jenner.ac.uk/VaxiJen) [28] and Allergen FP v.1.0 (http://ddg-pharmfac.net/AllergenFP) [29]. Moreover, the Discontinuous Bcell epitopes were also predicted using Disco tope 2.0 webserver tool [30] at a threshold value of −6.6 followed by representing the identified epitopes in the three-dimensional structure of Nsp13 protein. The Discotope score of residues above the threshold value indicates positive prediction and that below the threshold value indicates negative prediction, respectively.

## RESULTS

### Identification of mutations in Nsp13 among Indian SARS-Cov-2 isolates

The Nsp13 SARS-CoV-2 sequences reported from India till 24 May 2021 were aligned with the reference sequence reported from China (accession number: YP_009724389) using Clustal omega mediated multiple sequence alignments for mutational analysis. The amino acid variations in the Nsp13 protein sequences among Indian isolates were identified carefully and noted for analysis. The analysis revealed the presence of 38 mutations distributed all over the Nsp13 proteins and the details of each mutation are listed in the Table1 and their locations observed is represented in the three-dimensional structure of Nsp13 (PDB ID-6ZSL) using Pymol visualisation tool (Figure1). Our data also demonstrates the change in polarity and charge upon amino acids mutations in Nsp13. It was found that most of the mutations do not contribute to any change (Neutral to Neutral). Although, some of mutations led to alteration in charge from Basic to neutral (H164Y, H290Y, R392C and R442Q) while only one mutation at position 142 (H142V) altered the charge from acidic to neutral as depicted in the Table1.

**Figure 1:**
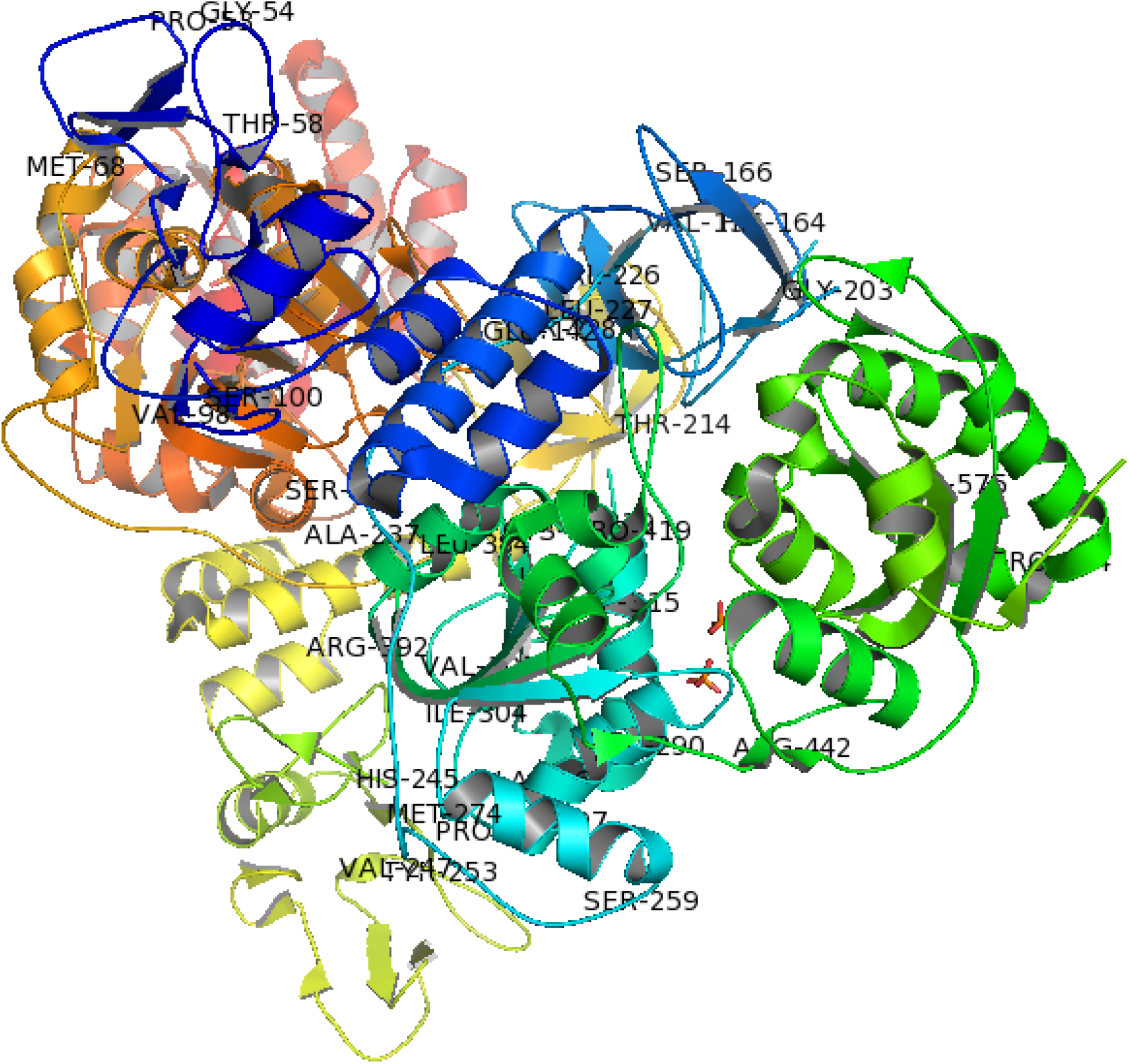
Three-dimensional structure of SARS-CoV-2 Nsp13 (PDB ID: 6ZSL) depicting the location of identified mutations in this study.

### Effect of mutation on stability and flexibility

The mutational analysis for evaluating the impact of mutations on the stability and flexibility of Nsp13 protein was conducted by DynaMut webserver. This webserver calculated the difference in free energy(ΔΔG) between wild type and mutant. The positive ΔΔG corresponds to increase in stability while the negative ΔΔG corresponds to decrease in stability. Our data demonstrate that a total of 38 mutations caused increase or decrease in the stability of Nsp13 protein. The positive ΔΔG values in the range of 0.013 to 0.975 kcal/mol shows stabilising effect (Table1). We have observed the maximum positive ΔΔG for H164Y (0.975), A237T (0.924), T214I (0.881), C309Y (0.844), S236I (0.750) and P419S (0.693), indicating stabilising mutations (Figure 2). While, V305 (−2.53), G54S (−1.69), A308Y (−1.306) shows the high negative ΔΔG value, which indicates the destabilization effect on the Nsp13 protein. (Figure 2). Similarly, the flexibility was also calculated by difference in vibrational entropy (ΔΔS) between wild type and mutant protein of Nsp13. The maximum positive ΔΔS (0.896) for H290Y caused increase in the flexibility of Nsp13 protein whereas, the maximum negative ΔΔS (−0.725) for A308Y represents the decrease in flexibility of protein (Table 2). Collectively, our data suggests that mutations identified in the Nsp13 Proteins could alter its stability and dynamicity.

**Table 2.**
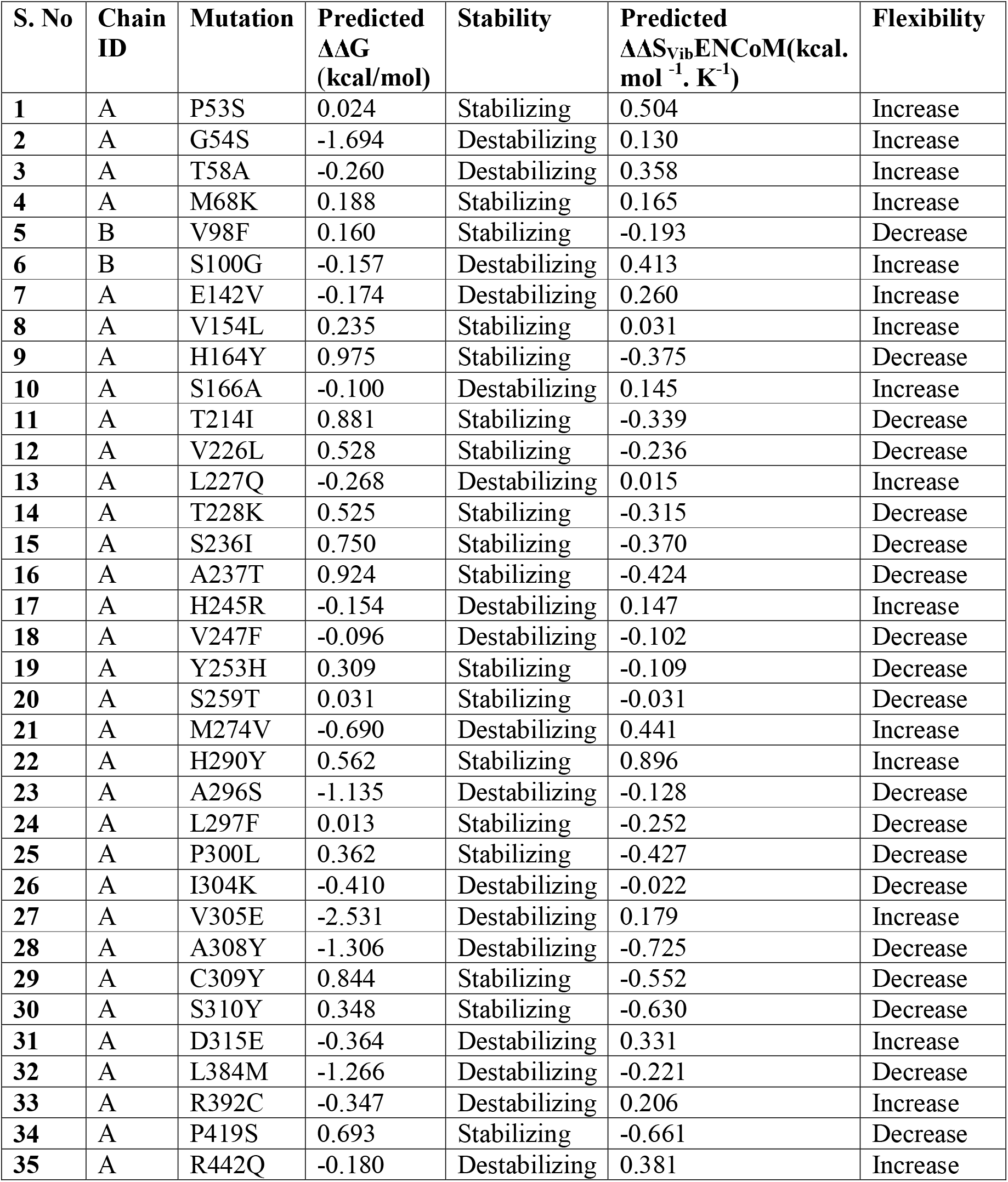
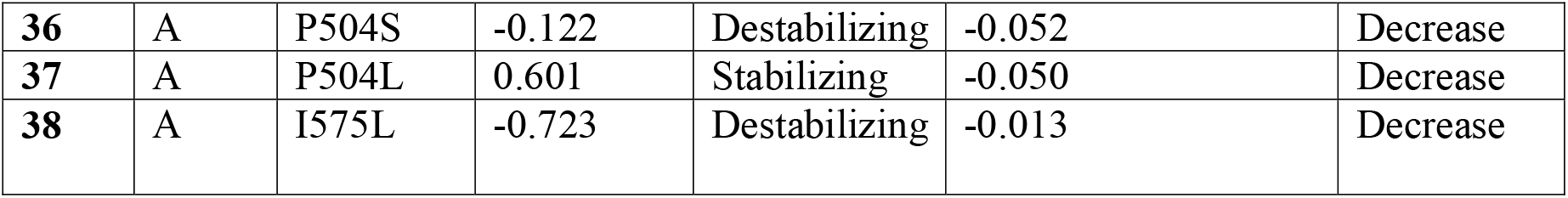
Predicted ΔΔG and ΔΔS_Vib_ between wild type and mutant Nsp 13 Protein.

**Figure 2:**
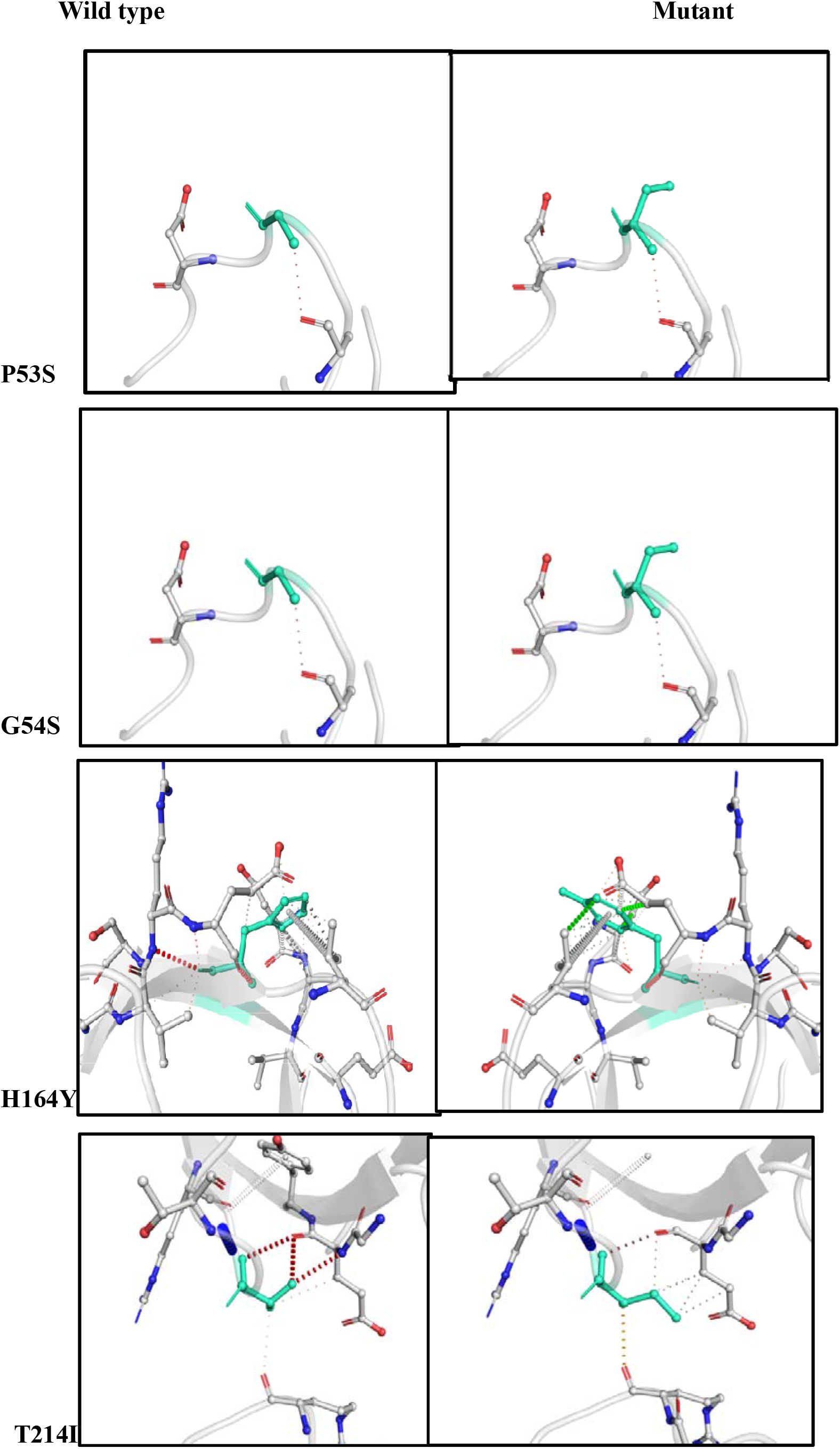
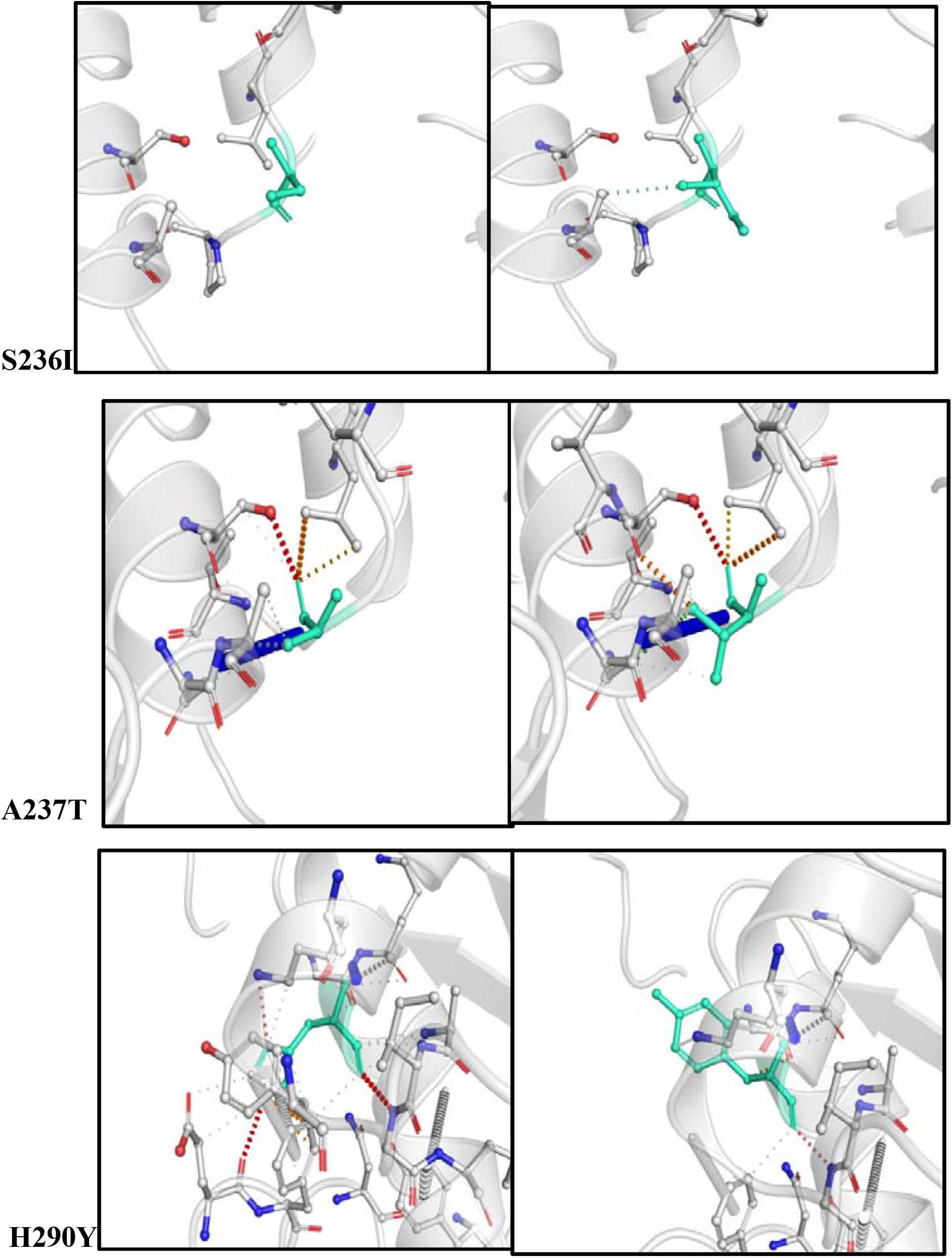
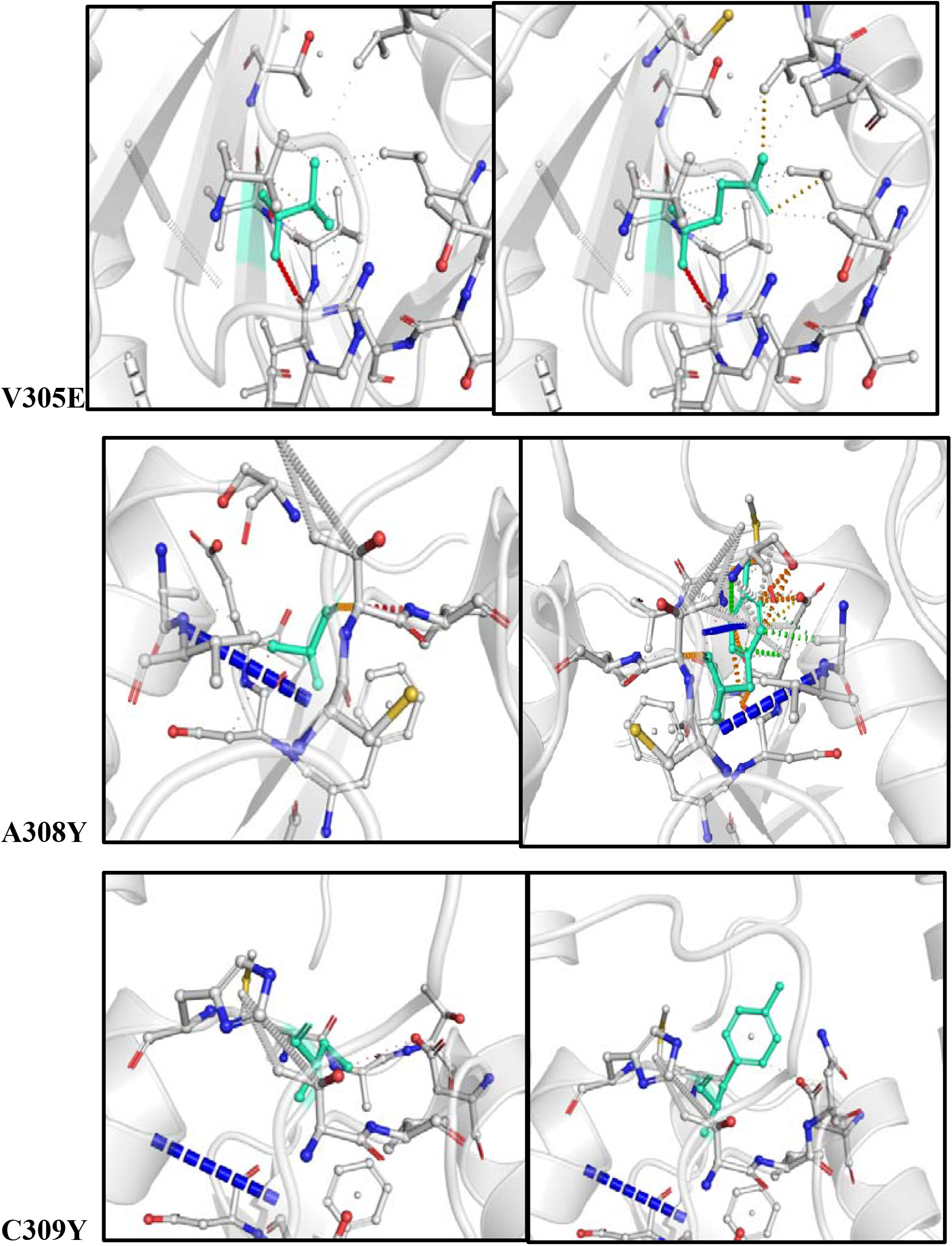
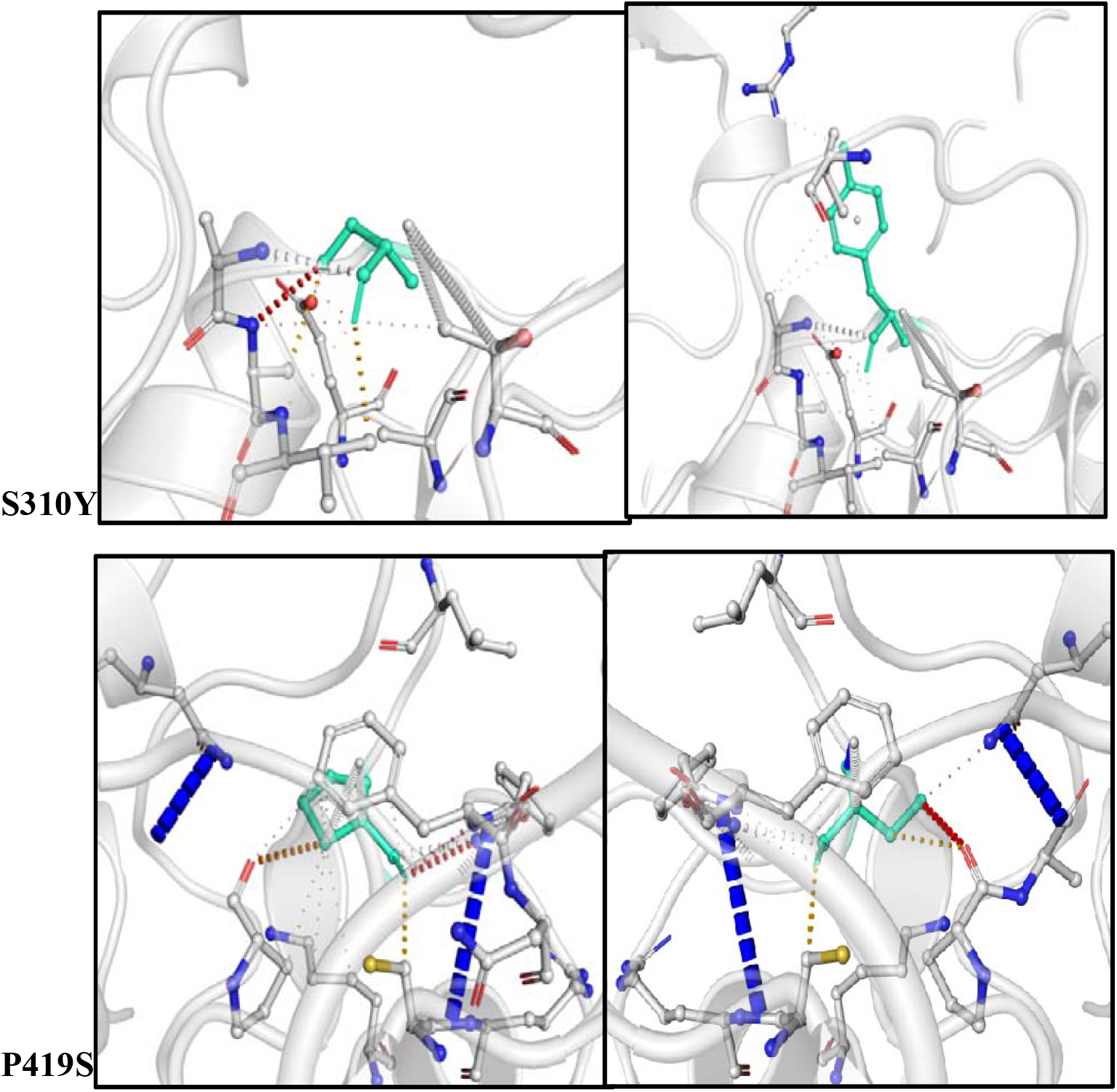
Visual representation of top 12 mutations contributed by Nsp13 that includes P53S,G54S, H164Y, T214I, S236I, A237T, C309Y, H290Y, V305E, A308Y, C309Y, S310Y and P419S, respectively. In each panel, the wild type and mutant residues of Nsp13 in their respective three dimensional structures are highlighted in the light green color. The intramolecular interactions between the highlighted residues and neighbouring residues in the Nsp13 protein are also indicated.

### Effect of Nsp13 mutation on protein function

The PROVEAN score predicts the functional impact of mutations in Nsp13. PROVEAN Score of mutants equal to or less than the default threshold score (−2.5) were predicted to have a deleterious effect on Protein whereas, PROVEAN Score more than the threshold score were predicted to have a neutral effect on the respective protein. A total of 38 mutations reported in Nsp13 among Indian SARS-CoV-2 sequences, includes 15 mutants as neutral and 23 mutants as deleterious based on their predicted PROVEAN Scores as mentioned in the Table 3.

**Table 3.** Predicted PROVEAN score of Nsp13 mutations and its impact on protein function.

| **S. No** | **Mutations** | **PROVEAN Score** | **Effect on protein** |
| --- | --- | --- | --- |
| 1 | P53S | -1.633 | Neutral |
| 2 | G54S | -4.046 | Deleterious |
| 3 | T58A | -0.506 | Neutral |
| 4 | M68K | -4.499 | Deleterious |
| 5 | V98F | -2.368 | Neutral |
| 6 | S100G | -3.933 | Deleterious |
| 7 | E142V | -6.883 | Deleterious |
| 8 | V154L | -0.076 | Neutral |
| 9 | H164Y | 1.312 | Neutral |
| 10 | S166A | -1.205 | Neutral |
| 11 | T214I | -5.162 | Deleterious |
| 12 | V226L | -2.763 | Deleterious |
| 13 | L227Q | -5.900 | Deleterious |
| 14 | T228K | -5.300 | Deleterious |
| 15 | S236I | -3.281 | Deleterious |
| 16 | A237T | -3.781 | Deleterious |
| 17 | H245R | 1.779 | Neutral |
| 18 | V247F | -1.795 | Neutral |
| 19 | Y253H | -1.608 | Neutral |
| 20 | S259T | -1.184 | Neutral |
| 21 | M274V | -1.983 | Neutral |
| 22 | H290Y | -5.300 | Deleterious |
| 23 | A296S | -1.750 | Neutral |
| 24 | L297F | -2.686 | Deleterious |
| 25 | P300L | -6.932 | Deleterious |
| 26 | I304K | -6.272 | Deleterious |
| 27 | V305E | -5.552 | Deleterious |
| 28 | A308Y | -5.900 | Deleterious |
| 29 | C309Y | -8.317 | Deleterious |
| 30 | S310Y | -5.786 | Deleterious |
| 31 | D315E | -3.933 | Deleterious |
| 32 | L384M | -1.983 | Neutral |
| 33 | R392C | -2.731 | Deleterious |
| 34 | P419S | -8.000 | Deleterious |
| 35 | R442Q | 0.467 | Neutral |
| 36 | P504S | -6.025 | Deleterious |
| 37 | P504L | -8.158 | Deleterious |
| 38 | I575L | -1.800 | Neutral |

### Prediction of B-cell epitopes

Linear continuous B-cell epitopes of SARS-CoV-2 Nsp13 were predicted by IEDB webserver tool as shown in figure 3A. The Bepipred score of Nsp13 residues depicts with a minimum score of −0.161 and a maximum score of −0.651 at threshold value of 0.482. The data set identified the top seven linear epitopes with at least eight amino acid residues, of which both their sequences and locations are mentioned in the Figure 3B. These epitopes were further analyzed for antigenicity, allergenicity and toxicity by *in-silico* based approaches. The predicted data shows that all of the seven peptides are non-toxic based on Toxin Pred prediction tool. Out of the seven peptides, three peptides are identified as allergen and four are non-allergen based on allergenicity prediction (Figure 3B). Similarly, two of these predicted linear epitopes exhibited antigenicity and the remaining five peptides as non-antigenic as analyzed by VaxiJen prediction tool (Figure 3B). In addition, discontinuous B-cell epitopes of Nsp13 were predicted using Discotope 2.0 server tool. The available 3D structure of Nsp13 in PDB format (PDB ID: 6ZSL) with chain A of protein used as input in the Discotope 2.0 server tool (Figure 3C). The residues with Discotope score equal or greater than threshold value (−6.6) was identified as discontinuous epitope residues of Nsp13. The data sets revealed the identification of 18 B cell epitope residues out of the total residues of Nsp13 protein at a threshold value of -6.6 (Figure 3D). Histidine and Valine at position 245 and 247, out of 18 predicted residues are significantly positively predicted with high Discotope score value as depicted in Figure 2D. Overall this data revealed B cell epitopes contributed by Nsp13.

**Figure 3:**
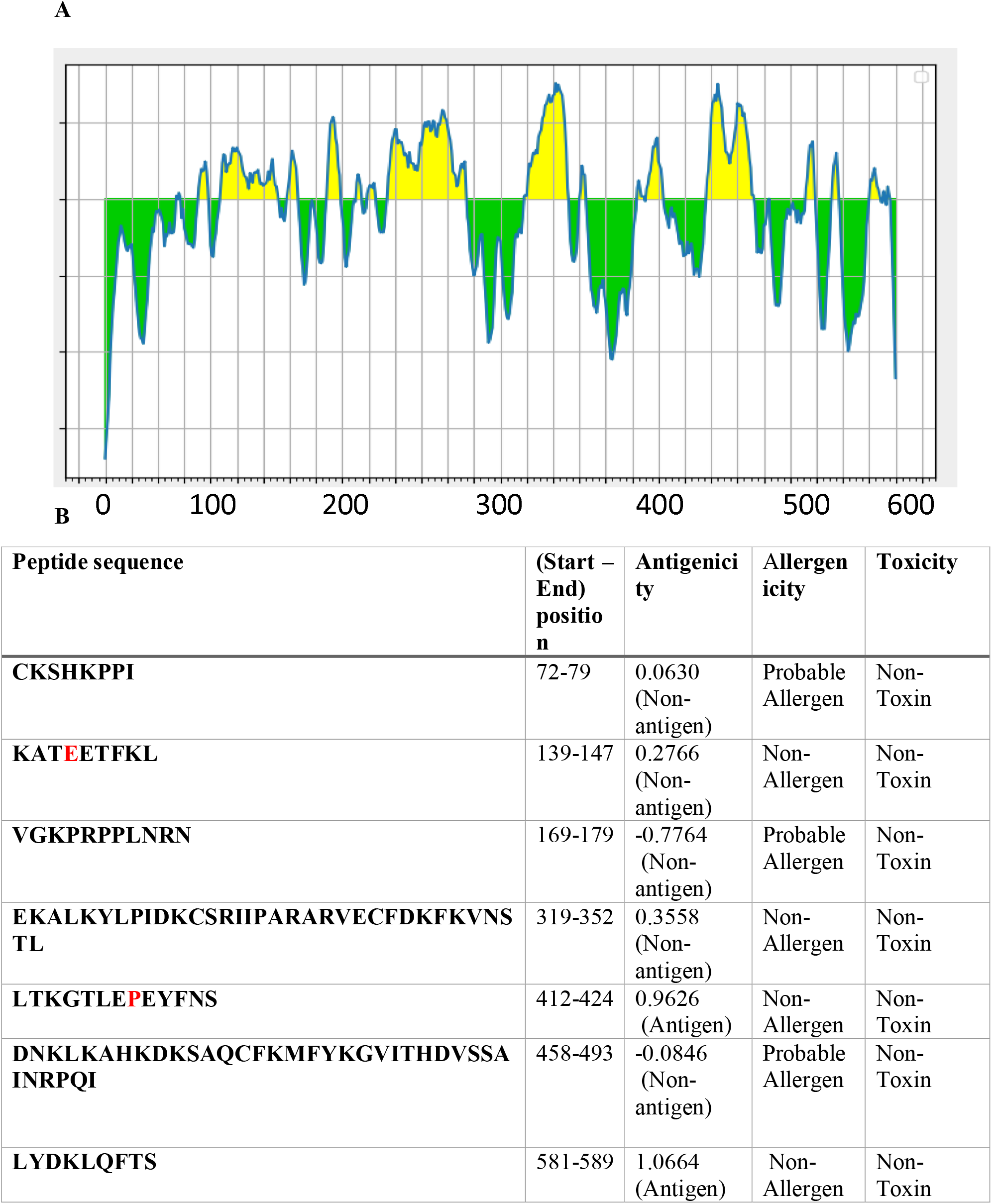
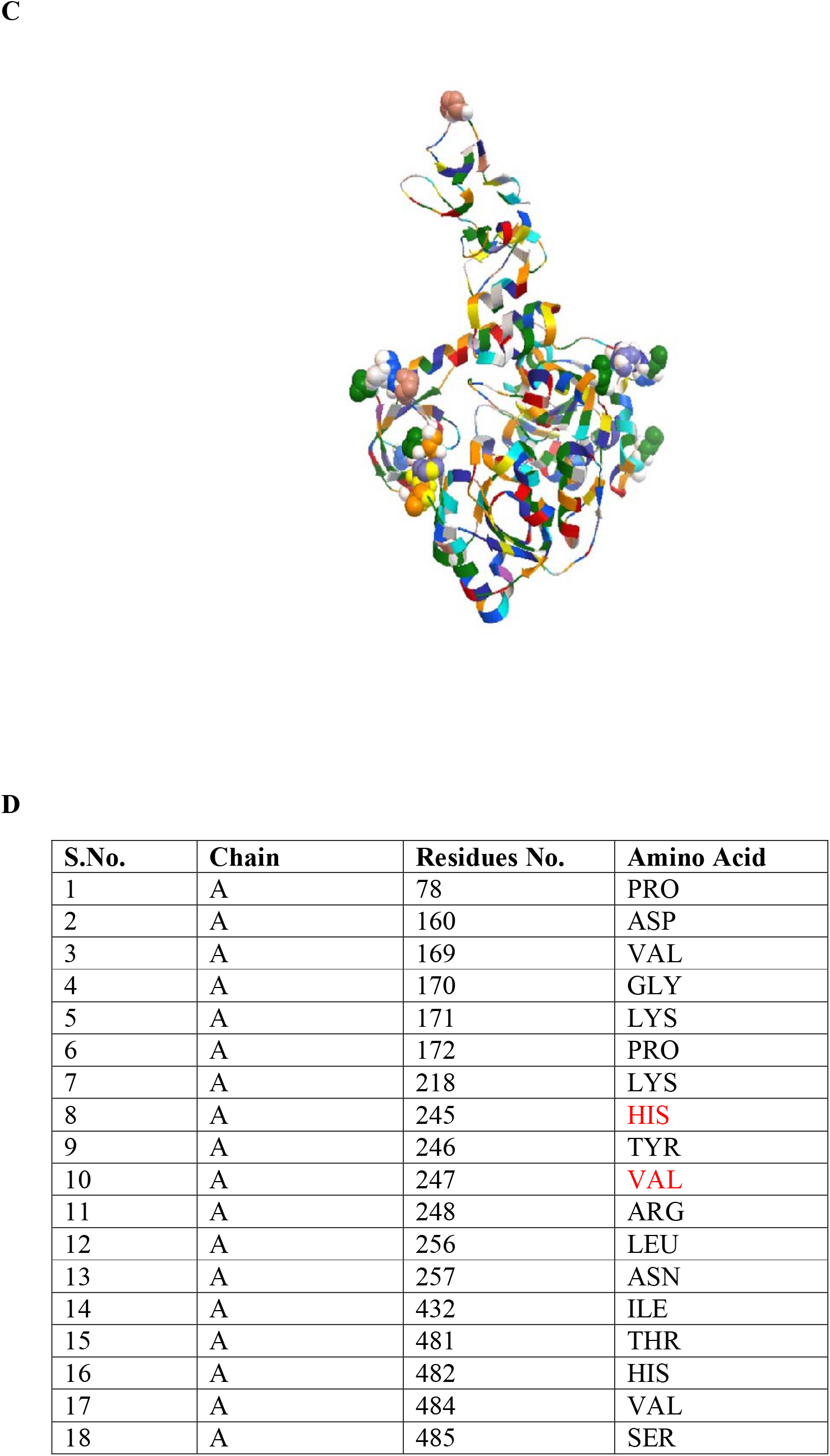
Prediction of B-cell epitopes of Nsp13. A) Prediction of linear B-cell epitopes. Y axis of the graph depicts the Bepipred Score, while the X-axis depicts the residues position of Nsp13. The yellow area of the graph indicates the part of protein with higher probability to be localised as epitope. B) The top seven predicted peptides of Nsp13. The red font color indicate the location of mutant residues in the peptide sequence. C) Prediction of Discontinuous B-cell epitopes. The prediction of discontinuous epitope residues on the 3-D structure of Nsp13 (PDB ID: 6ZSL). D) The position and identity of each discontinuous epitope residues of Nsp13.

## DISCUSSIONS

In the present study, we identified 38 mutations in Nsp13 from Indian isolates. The variations in the SARS-CoV-2 proteins enable us to better understand its diversity, dynamics and genetic epidemiology [31] that might provide an opportunity in developing effective and safe therapeutic drugs and vaccines for this highly mutable coronavirus. Our data revealed that the observed mutations at different positions (H164Y, A237T, T214I, C309Y, S236I, P419S, V305E, G54S, H290Y, P53S, A308Y and A308Y) in the Nsp13 significantly affect the stability and flexibility of Nsp13 protein. In addition, the alterations in the function of Nsp13 were also predicted upon mutational effect by their Predicted PROVEAN scores. Furthermore, B-cell epitopes contributed by Nsp13 were identified using various predictive immunoinformatic tools. Immunological Parameters of Nsp13 such as antigenicity, allergenicity and toxicity were assessed to predict the potential B-cell epitopes. Predicted *in-silico* epitopes are under urgent consideration in the development of epitope-based peptide vaccine candidates against SARS-CoV-2. The predicted peptide sequences were correlated with the observed mutants. Our predicted data showed that there are seven high rank linear epitopes as well as 18 discontinuous B-cell epitopes based on immunoinformatic findings. Moreover, it was investigated that, out of total 38 identified mutations among Indian SARS-CoV-2 Nsp13 protein, four mutant residues at position 142 (E142), 245 (H245), 247 (V247) and 419 (P419) are localised in the predicted B cell epitopic region (Figure 3). These mutant residues may help the SARS-CoV-2 variants in eliciting a distinct immune response from the wild type SARS-CoV-2. Several previous studies evidently support the findings with similar observations using the immunoinformatic approaches [32-34]. Hence, our study provides some insights in understanding the Nsp13 epitopes which could regulate host immune responses against SARS-CoV-2. The present *in-silico* study revealed the impact of identified mutations on the stability and flexibility of Nsp13 protein. Further *in-vivo* studies are necessary to better understand and validate the effect of mutations on the immunogenicity of epitopes of SARS-CoV-2.

## CONCLUSION

The emergence of a novel coronavirus, SARS-CoV-2 has become a major global concern with an unprecedented public health crisis. Understanding the genomic variations in SARS-CoV-2 could help in improving the current diagnostic techniques and developing suitable vaccine candidates against the SARS-CoV-2 infections. Altogether, the results of the present *in-silico* study show the various mutations in Nsp13 from Indian SARS-CoV-2 and their implications on stability, flexibility and function of Nsp13 protein using predictive tools of immunoinformatics. Non-structural proteins exhibit low glycation density as compared to the structural proteins of SARS-CoV-2. Therefore, epitopes of Nsp13 could be used as effective and promising target against SARS-CoV-2.

## Author contributions

AK has conceived the study and contributed in study-design, data interpretation, and manuscript preparation. DK has performed the experiments, analyzed data, and prepared manuscript. NK, SK, PKS, SKS and NRB have contributed in the conceptualization of the study and reviewing the manuscript.

## ACKNOWLEDGEMENTS

Declare none

---

Supplementary Table 1: list of protein accession number used in the study.

