## supplementary Table 1 for "Identification and characterization of novel mutants of Nsp13 protein among Indian SARS-CoV-2 isolates"

Supplementary table1: list of protein accession number used in the study.

| Yp_009724389 | QVO44046 | QVL01198 | QRN68424 | QPB18158 | QOS50688 | QOQ57034 |
| --- | --- | --- | --- | --- | --- | --- |
| QVR42238 | QVO44058 | QVI17109 | QRN68442 | QPB18194 | QOS50712 | QOQ57046 |
| QVR42250 | QVO44070 | QVI17121 | QRM36235 | QPB18206 | QOS50724 | QOQ57058 |
| QVR43916 | QVO44086 | QTV76173 | QRM91443 | QPB39980 | QOS50736 | QOQ57070 |
| QVR43928 | QVO44098 | QTP17383 | QRE02629 | QPB39992 | QOS50748 | QOQ57082 |
| QVR43940 | QVO44110 | QTH36343 | QRC50281 | QPB40004 | QOS50760 | QOQ57094 |
| QVR43952 | QVO44133 | QTH36355 | QRC50293 | QPA18453 | QOS50772 | QOQ57106 |
| QVR43964 | QVO44145 | QTH36367 | QRC50305 | QPA18457 | QOS50784 | QOQ57119 |
| QVR48690 | QVO44206 | QTH36379 | QRC50317 | QPA18459 | QOS50796 | QOQ72542 |
| QVR52999 | QVO59419 | QTH36391 | QRC50325 | QOW08181 | QOS50808 | QOQ72554 |
| QVR83663 | QVP26002 | QTH36403 | QRC50340 | QOU99120 | QOS50820 | QOQ72566 |
| QVR83675 | QVP26014 | QTH36415 | QRC50352 | QOU99144 | QOS50843 | QOQ72578 |
| QVS02345 | QVP26026 | QTH36427 | QRC50364 | QOU99168 | QOS50855 | QOQ84793 |
| QVS02357 | QVP26038 | QTH36439 | QRC50376 | QOU99180 | QOS50867 | QOQ84824 |
| QVS02369 | QVP26050 | QTH36451 | QRC50492 | QOU99192 | QOS50879 | QKM77202 |
| QVS03027 | QVP26064 | QTH36723 | QQJ94103 | QOU99203 | QOS50891 | QKM77214 |
| QVS16977 | QVP26076 | QTG68689 | QQJ95306 | QOU99214 | QOS50926 | QKM77226 |
| QVS16989 | QVP26088 | QTG68602 | QQH18637 | QOU99225 | QOS50938 | QKM77238 |
| QVS30945 | QVP26100 | QTG68615 | QPZ33508 | QOU99237 | QOS50950 | QKM77250 |
| QVS30957 | QVP26118 | QTG40603 | QPB40017 | QOU99249 | QOS50962 | QKM77262 |
| QVS48980 | QVP26130 | QRY06633 | QPB40029 | QOU99261 | QOS50974 | QKM77274 |
| QVS48992 | QVP26142 | QRY06645 | QPB40041 | QOU99272 | QOS50986 | QKM77286 |
| QVS53168 | QVP26154 | QRY06657 | QPB40053 | QOU99283 | QOS51010 | QNO30921 |
| QVR41769 | QVP26486 | QRY06669 | QPB40065 | QOU99294 | QOS51022 | QNO30933 |
| QVR41781 | QVP26501 | QRY06681 | QPB40077 | QOS50439 | QOS51034 | QNN87972 |
| QVQ64917 | QVP26513 | QRQ47003 | QPB40089 | QOS50449 | QOS51046 | QNN87984 |
| QVQ64929 | QVP26525 | QRQ47018 | QPB40101 | QOS50461 | QOS51058 | QNN87996 |
| QVQ64943 | QVP30914 | QRQ47032 | QPB40113 | QOS50473 | QOS51070 | QNN88008 |
| QVQ64955 | QVP43688 | QRQ47071 | QPB40563 | QOS50497 | QOS51082 | QNN88020 |
| QVQ64967 | QVP43700 | QRQ47083 | QPB40575 | QOS50509 | QOS51094 | QNN88068 |
| QVQ64981 | QVP57055 | QRQ69195 | QPB17918 | QOS50521 | QOS51106 | QNN88080 |
| QVQ64993 | QVP61496 | QRQ69279 | QPB17990 | QOS50545 | QOS51118 | QNN88092 |
| QVQ65005 | QVP61508 | QRN68229 | QPB18014 | QOS50557 | QOR63432 | QNN88104 |
| QVQ65017 | QVP79437 | QRN68241 | QPB18026 | QOS50580 | QOR63444 | QNN88116 |
| QVQ65029 | QVP88490 | QRN68253 | QPB18050 | QOS50592 | QOR63456 | QNN88128 |
| QVO43863 | QVP88502 | QRN68266 | QPB18062 | QOS50604 | QOR63468 | QNN88140 |
| QVO43969 | QVQ02196 | QRN68278 | QPB18074 | QOS50616 | QOR63480 | QNN88152 |
| QVO43981 | QVQ15054 | QRN68290 | QPB18086 | QOS50628 | QOR63504 | QNN88164 |
| QVO43993 | QVQ15066 | QRN68302 | QPB18098 | QOS50640 | QOR64231 | QNN88176 |
| QVO44011 | QVQ19425 | QRN68315 | QPB18110 | QOS50652 | QOR64243 | QNN88188 |
| QVO44023 | QVL01149 | QRN68400 | QPB18122 | QOS50664 | QOQ57010 | QNN88200 |
| QVO44034 | QVL01186 | QRN68412 | QPB18134 | QOS50676 | QOQ57022 | QNN88212 |

| QNN88224 | QNL90822 | QLQ87442 | QLR12259 | QLH90076 | QLA09820 | QKY60223 |
| --- | --- | --- | --- | --- | --- | --- |
| QNN88236 | QNL90834 | QLQ87454 | QLR12271 | QLH90088 | QLA09832 | QKY60235 |
| QNN88248 | QNL90846 | QLQ87466 | QLR12283 | QLH90101 | QLA09844 | QKY60262 |
| QNN88260 | QNL90858 | QLQ87478 | QLR12295 | QLH93117 | QLA09856 | QKY64328 |
| QNN88272 | QNL90870 | QLQ87490 | QLR12307 | QLH93129 | QLA09880 | QKY64344 |
| QNN88284 | QNL90882 | QLQ87502 | QLR12319 | QLH93185 | QLA09904 | QKY64358 |
| QNN88296 | QNL90894 | QLQ87514 | QLR12331 | QLH93199 | QLA10066 | QKY64612 |
| QNN88551 | QNL90906 | QLQ87526 | QLR12343 | QLH93212 | QLA10078 | QKY64638 |
| QNN88563 | QNL90918 | QLQ87550 | QLR12355 | QLH93283 | QLA10090 | QKY64790 |
| QNN88575 | QNL98443 | QLQ87574 | QLR12415 | QLF97937 | QLA10102 | QKY65275 |
| QNN88587 | QNL98455 | QLQ87598 | QLR12438 | QLF97949 | QLA10114 | QKV25883 |
| QNN88599 | QNL98467 | QLQ87610 | QLR60574 | QLF97985 | QLA10126 | QKV25895 |
| QNN88630 | QNL98479 | QLQ87622 | QLR80398 | QLF97997 | QLA10138 | QKV25907 |
| QNN88642 | QNL35816 | QLQ87634 | QLS04049 | QLF98069 | QLA10150 | QKV25919 |
| QNN88654 | QNL35828 | QLQ87658 | QLI49695 | QLF98081 | QLA10162 | QKV25931 |
| QNN90122 | QNL35840 | QLQ87682 | QLI49707 | QLF98093 | QLA10174 | QKV25943 |
| QNN90135 | QNL35852 | QLQ87694 | QLI49719 | QLF98105 | QLA10186 | QKV25955 |
| QNN93013 | QNL35864 | QLQ87706 | QLI49731 | QLF98117 | QLA10198 | QKV25967 |
| QNN26334 | QNL35876 | QLQ87718 | QLI49743 | QLF98129 | QLA10210 | QKV25979 |
| QNN26346 | QNL35888 | QLQ87730 | QLI49755 | QLF98141 | QLA10222 | QKV25991 |
| QNN26358 | QNL35900 | QLQ87795 | QLI49767 | QLF98153 | QKY74626 | QKV26003 |
| QNN26370 | QNL35912 | QLQ87983 | QLI49779 | QLF98165 | QKY74638 | QKV26015 |
| QNN26382 | QNL35924 | QLQ87995 | QLI49791 | QLF98178 | QKY59939 | QKV26027 |
| QNN26394 | QNL35936 | QLQ88046 | QLI49803 | QLF98198 | QKY59951 | QKV26039 |
| QNN26406 | QNL35948 | QLQ88063 | QLI49815 | QLF98210 | QKY59963 | QKV26051 |
| QNN26418 | QNL35960 | QLQ88075 | QLI52055 | QLF98222 | QKY59975 | QKV26063 |
| QNN26430 | QNL35972 | QLQ88087 | QLI52067 | QLF98234 | QKY59987 | QKV26075 |
| QNN26442 | QNL35984 | QLQ88102 | QLH64777 | QLF98246 | QKY59999 | QKV26087 |
| QNN26454 | QNL35996 | QLR06736 | QLH64789 | QLF98258 | QKY60023 | QKV27549 |
| QNN26466 | QNL36008 | QLR06757 | QLH64801 | QLF98275 | QKY60047 | QKV27561 |
| QNN26478 | QNL36560 | QLR06769 | QLH64813 | QLF98287 | QKY60059 | QKV27573 |
| QNN30834 | QIA98563 | QLR06870 | QLH64825 | QLA09686 | QKY60071 | QKV27585 |
| QNN30858 | QIA98573 | QLR06882 | QLH64837 | QLA09698 | QKY60083 | QKQ29884 |
| QNN30870 | QLY82444 | QLR06894 | QLH64849 | QLA09710 | QKY60095 | QKQ29944 |
| QNN31194 | QLY82482 | QLR07146 | QLH64861 | QLA09722 | QKY60107 | QKQ29956 |
| QNN31206 | QLY82500 | QLR07158 | QLH64875 | QLA09734 | QKY60119 | QKQ29968 |
| QNN81330 | QLY82512 | QLR07170 | QLH64887 | QLA09746 | QKY60131 | QKQ30028 |
| QNN81490 | QMC85346 | QLR07182 | QLH64899 | QLA09758 | QKY60163 | QKQ30040 |
| QNN81502 | QLQ87394 | QLR07194 | QLH64915 | QLA09770 | QKY60175 | QKQ30052 |
| QNN83660 | QLQ87406 | QLR07206 | QLH64927 | QLA09782 | QKY60187 | QKQ30064 |
| QNN83672 | QLQ87418 | QLR12235 | QLH64939 | QLA09794 | QKY60199 | QKQ30076 |
| QNL90810 | QLQ87430 | QLR12247 | QLH90040 | QLA09808 | QKY60211 | QKQ30088 |

| QKQ30100 | QKJ68723 | QJY77053 | QJX44392 | QJT43546 | QJQ28355 |
| --- | --- | --- | --- | --- | --- |
| QKQ30112 | QKJ68735 | QJY51262 | QJX44404 | QJT43558 | QJQ28367 |
| QKQ30124 | QKJ84941 | QJY51274 | QJX44416 | QJT43570 | QJQ28379 |
| QKQ30136 | QKJ84953 | QJY51286 | QJX44428 | QJT43582 | QJQ28391 |
| QKQ30148 | QKJ84965 | QJY51346 | QJX44440 | QJT43594 | QJQ28403 |
| QKQ30160 | QKJ84977 | QJY51358 | QJX44464 | QJT43606 | QJQ28415 |
| QKQ30172 | QKG91164 | QJY51370 | QJX44476 | QJT43618 | QJQ28427 |
| QKQ30184 | QKG91176 | QJY51382 | QJX44500 | QJT43630 | QJH92165 |
| QKQ30196 | QKG91188 | QJY40383 | QJX44512 | QJT43642 | QJH92177 |
| QKQ30208 | QKG91200 | QJY40395 | QJX44536 | QJT43654 | QJF77844 |
| QKQ30220 | QKG91212 | QJY40407 | QJX44548 | QJT43666 | QJF77856 |
| QKQ30232 | QKG91224 | QJY40419 | QJX44560 | QJT43678 | QJF77868 |
| QKQ30244 | QKG91236 | QJY40431 | QJX44572 | QJT43690 | QJF77880 |
| QKQ30256 | QKG91248 | QJY40443 | QJX44584 | QJT43702 | QJF11810 |
| QKQ30268 | QKG91260 | QJY40455 | QJX44596 | QJT43714 | QJF11822 |
| QKQ63386 | QKG91272 | QJY40467 | QJX44608 | QJT43726 | QJF11834 |
| QKO00484 | QKG91284 | QJY40479 | QJX44620 | QJS39637 | QJF11846 |
| QKJ68410 | QKH78808 | QJY40491 | QJX44632 | QJS39649 | QJF11858 |
| QKJ68422 | QKH78820 | QJY40503 | QJX44644 | QJR84343 | QJF11870 |
| QKJ68434 | QKI10470 | QJY40515 | QJX44656 | QJR84355 | QJF11882 |
| QKJ68446 | QKI28575 | QJY40527 | QJX44668 | QJR84367 | QJC19489 |
| QKJ68458 | QKI28587 | QJY40539 | QJX44680 | QJR84379 | QHS34545 |
| QKJ68471 | QKI28599 | QJY40551 | QJW39842 | QJR84391 | QIA98582 |
| QKJ68483 | QKI28611 | QJY40563 | QJW39854 | QJR84403 |  |
| QKJ68495 | QKI28623 | QJY40575 | QJW39866 | QJR84415 |  |
| QKJ68507 | QKI28635 | QJY40587 | QJW39878 | QJR84427 |  |
| QKJ68519 | QKI28647 | QJY40599 | QJW39890 | QJR84439 |  |
| QKJ68531 | QKI28659 | QJW00289 | QJW39902 | QJR84451 |  |
| QKJ68543 | QKI28671 | QJW00301 | QJW39914 | QJR84463 |  |
| QKJ68555 | QKI28683 | QJW00313 | QJW39926 | QJR84475 |  |
| QKJ68567 | QKE61658 | QJW00325 | QJW69137 | QJR84487 |  |
| QKJ68579 | QKE61670 | QJW00337 | QJU70555 | QJR84499 |  |
| QKJ68591 | QKE61682 | QJW00349 | QJT43414 | QJR84511 |  |
| QKJ68603 | QKE61694 | QJW00361 | QJT43426 | QJR84523 |  |
| QKJ68615 | QKE61706 | QJW00373 | QJT43438 | QJR84535 |  |
| QKJ68627 | QKE61718 | QJW00385 | QJT43450 | QJQ39966 |  |
| QKJ68639 | QKE61730 | QJW00397 | QJT43462 | QJQ39978 |  |
| QKJ68651 | QKE61742 | QJW00409 | QJT43474 | QJQ39990 |  |
| QKJ68663 | QKE61754 | QJW00421 | QJT43486 | QJQ27840 |  |
| QKJ68675 | QKE61766 | QJW00433 | QJT43498 | QJQ27852 |  |
| QKJ68687 | QKE61778 | QJW00445 | QJT43510 | QJQ27864 |  |
| QKJ68699 | QKE61790 | QJW00457 | QJT43522 | QJQ27876 |  |
| QKJ68711 | QKE61802 | QJX44380 | QJT43534 | QJQ28343 |  |
